# SNIPE score can capture prodromal Alzheimer’s in cognitively normal subjects

**DOI:** 10.1101/541854

**Authors:** Azar Zandifar, Vladimir Fonov, Olivier Potvin, Simon Duchesne, D. Louis Collins, for the Alzheimer’s Disease Neuroimaging Initiative

## Abstract

Capturing early changes in the brain related to Alzheimer’s disease may lead to models that successfully predict cognitive decline and the eventual onset of dementia, well ahead of onset of clinical symptoms. In this study we used both hippocampal volume and our hippocampal driven SNIPE score to show which marker better captures Alzheimer’s related changes in a large dataset of normal controls (N=515) from the ADNI study, comparing controls that remain cognitively stable and controls that progress to either MCI or Alzheimer’s dementia during 10 years of follow-up (median follow-up: 30 months). We measured hippocampal volume and SNIPE score and found that the effect size to differentiate between stable and progressor groups was significantly larger for SNIPE score than for volume. Our results also show that there is a significant age-related difference between groups for both markers, and the difference is greater with the SNIPE score. Our experiments show that considering high sensitivity of our SNIPE score regarding to early AD-related brain changes, this marker is a better candidate in comparison to hippocampal volume for predicting the future onset of dementia.

## Introduction

Alzheimer’s disease (AD) is characterized by abnormal tau aggregation, concurrent to synaptic dysfunction, cell death, and brain atrophy (Oddo et al., 2006), as well as abnormal processing of the amyloid precursor protein, which leads to beta amyloid deposits in the cortex (Blennow, de Leon, & Zetterberg, 2006). In its typical non-dominantly inherited form (>99% of cases)(Campion et al., 1999), sufficient evidence has been gathered from biomarker studies (C. R. Jack, Jr. et al., 2013) as well as post-mortem, pathological reports (Duyckaerts, 2011; Hyman et al., 2012) to postulate that these processes span more than two decades. Given such evidence, early detection of prodromal disease is key to intervene before the onset of too much irreversible neurodegeneration. This requires biomarkers specific to AD progression that are sufficiently sensitive decades ahead of diagnostic.

Radiological-pathological studies have confirmed that structural (T1-weighted) MRI tracks brain atrophy in AD (Csernansky et al., 2004; C. R. Jack, Jr. et al., 2002) at the global, lobar and regional level (Ashburner et al., 2003; Chetelat & Baron, 2003; Csernansky et al., 2004; Fox & Schott, 2004), shown to correspond to neuronal losses in layer II of the entorhinal cortex (Gomez-Isla et al., 1996), in hippocampal CA1 (West, Coleman, Flood, & Troncoso, 1994), in the superior temporal gyrus (Gomez-Isla et al., 1997), and in the supramarginal gyrus (Grignon, Duyckaerts, Bennecib, & Hauw, 1998). Hippocampal atrophy in particular has been thoroughly studied in AD (H. Braak & Braak, 1991; Heiko Braak & Braak, 1995), with a clear reported difference between patients and age-matched cognitively healthy (CH) individuals (Coupé, Eskildsen, Manjón, Fonov, & Collins, 2012; C. R. Jack et al., 1997; A. Zandifar et al., 2017). In those subjects with mild cognitive impairment (MCI), that is those with memory complaints who objectively show demonstrable cognitive abnormalities and biomarker evidence of AD pathology but do not meet criteria for dementia (Albert et al., 2011), hippocampal volume can predict progression on a single subject basis (Coupé, Eskildsen, Manjón, Fonov, & Collins, 2012; A. Zandifar et al., 2017; Azar Zandifar, Fonov, Coupé, Pruessner, & Collins, 2014). It has been shown as well that hippocampal volume shows very low percentage of abnormality in cognitively healthy cohorts, while abnormality grows with disease progression (C. R. Jack et al., 2013). Our recent study shows that classification of AD patients versus CH based on hippocampal volume yields an area under the receiver operating curve (AUC) for the AD Neuroimaging Initiative (ADNI) dataset of 88%, while the AUC is 64% for differentiating MCI individuals that progressed to probable AD from those whose status remained stable for up to three years of follow-up, a result that did not depend on the different automatic segmentation methods tested (A. Zandifar et al., 2017). Hippocampal volume and atrophy stand therefore as putative biomarkers that could be used for early detection.

However, the Scoring by Nonlocal Image Patch Estimator (SNIPE) metric has been shown to surpass hippocampal volume in terms of predictive power, especially in the MCI stage (Coupé, Eskildsen, Manjón, Fonov, & Collins, 2012; Coupe et al., 2015). When compared to hippocampal volume, the SNIPE score increased the accuracy of a single-subject prediction of dementia in the MCI population by 10 percent (Coupé, Eskildsen, Manjón, Fonov, Pruessner, et al., 2012). Inspired by nonlocal patch-based denoising (Coupe et al., 2008), SNIPE is a disease probability scoring method, estimated within the hippocampal and entorhinal cortices (Coupé, Eskildsen, Manjón, Fonov, & Collins, 2012), which assigns a similarity score to each voxel that shows how much the patch around that specific voxel is similar to a library of probable AD patients or age-matched CH subjects.

Our aim in this study was to demonstrate the sensitivity of SNIPE scoring at detecting very early AD-related pathological changes in a cognitively healthy cohort. Our hypothesis was that there would be a difference in SNIPE scores between CH individuals that remained cognitively stable from those who declined, well before the onset of clinical symptoms; and that this difference was more emphasized using hippocampal SNIPE scoring than hippocampal volumetry.

## Methods

### Participants and clinical follow-up

Data used in the preparation of this article were obtained from the Alzheimer’s Disease Neuroimaging Initiative (ADNI) database (adni.loni.usc.edu). The ADNI was launched in 2003 as a public-private partnership, led by Principal Investigator Michael W. Weiner, MD. The primary goal of ADNI has been to test whether serial magnetic resonance imaging (MRI), positron emission tomography (PET), other biological markers, and clinical and neuropsychological assessment can be combined to measure the progression of mild cognitive impairment (MCI) and early Alzheimer’s disease (AD). For up-to-date information, see http://www.adni-info.org.

The dataset used in this study is composed of the CH cohort of ADNI-1 and ADNI-2. They showed no signs of depression, mild cognitive impairment or dementia (Alzheimer’s Disease Neuroimaging Initiative, 2018). The dataset consisted of 515 CH individuals, 228 from ADNI-1 and the balance from ADNI-2. Each subject was followed throughout the study period (up to 10 years) to screen for any kind of future change in diagnostic label. We used the most recent clinical diagnostic information available for each subject to determine if these individuals remained stable or progressed to either MCI or AD. From our dataset of 515 normal controls at baseline, 427 maintained their cognitively normal status throughout the follow-up period and were thus labeled as *Stables*. The remaining 88 subjects converted to either MCI or AD and were labeled as *Progressives*. The median follow-up period (30 months) is the same for the two groups. Study phase, age, and sex were extracted from baseline reports, and are summarized in Table 1.

**Table 1.** Dataset Information - The values in parentheses are 25% and 75% quantiles respectively.

|  | **Progressive** | **Stable** |
| --- | --- | --- |
| **Number** | 88 | 427 |
| **Median Age at Baseline** | 76.4 (72.5-80.0) | 73.8 (70.1-77.6) |
| **Sex: Female (%)** | 43 | 44 |
| **Median Education (yrs)** | 16 (14.75-18) | 16 (14-18) |
| **Median MMSE score** | 29 (29-30) | 29 (29-30) |
| **Median follow (months)** | 30 (24-55.5) | 30 (18-48) |

### Images and preprocessing pipeline

Baseline T1-weighted (T1w) magnetic resonance images (MRIs) were downloaded from ADNI and processed initially through a fully automatic pipeline (Aubert-Broche et al., 2013) that consisted in denoising (Coupe et al., 2008), correction of inhomogeneity using N3 (Sled, Zijdenbos, & Evans, 1998), registration to pseudo-Talairach stereotaxic space (Collins, Neelin, Peters, & Evans, 1994) using a population-specific template (Fonov et al., 2011), and brain extraction using BEaST (Eskildsen et al., 2012).

### Hippocampal Volumetry

The hippocampus was segmented automatically using a multi-template patch-based segmentation method (Coupé et al., 2011; A. Zandifar et al., 2017), which uses a training library for MRI volumes with manually traced hippocampi. While the patch-based strategy drastically increases the number of sample library patches involved in label assignment, it reduces susceptibility to registration error. The final label was assigned based on non-local means: each patch was weighted based on its similarity to the target patch and the final label (hippocampus or background) was assigned based on a weighted average over all similar patches. The hippocampal volume was estimated by counting voxels in pseudo-Talairach stereotaxic space, thus inherently normalized for head size.

### Hippocampal SNIPE Scoring

A similar approach to the segmentation strategy was followed to assign a grading value to each hippocampi (Coupé, Eskildsen, Manjón, Fonov, & Collins, 2012). The SNIPE score metric shows how much an image patch is intensity-wise similar to either a cognitively healthy or an AD cohort of training subjects. In fact, instead of hippocampal labels, the diagnosis label of the subjects in the template library is incorporated (−1 for AD, +1 for CH). The similarity for each voxel is defined based on its corresponding patch. The final score of each voxel is defined based on non-local average over the most similar patches.

The group similarity metric was averaged over the structure area, which is defined by the segmentation process, to assign a value to the whole structure (Coupé, Eskildsen, Manjón, Fonov, & Collins, 2012). In this study, we computed volumes and SNIPE scores for both left and right hippocampi on our whole cohort using the same training library as the one described in (Coupe et al., 2015). The template library images are all drawn from ADNI-1 dataset. There was no statistically significant difference between the AD and CN group in age or gender using a Generalized Linear Model (GLM). Since some of the cognitively healthy subjects are already included in the training library, when computing the SNIPE score for that subject, the corresponding MRI was removed from the training library.

### Statistical analyses and metrics

All statistical analyses were done using RStudio working under R 3.3.2.

To investigate the sensitivity of each biomarker in detecting between-group differences, we computed the Cohen’s d effect size based on both hippocampal volumes and hippocampal SNIPE scores, correcting for age, sex, and ADNI study phase. The correction is done using a similar method to the one presented in (Dukart, Schroeter, Mueller, & Initiative, 2011) for neurodegeneration and dementia. A linear regression is fitted using only normal controls to regress out the effect of confounding factors such as age while preserving the effect of atrophy-induced neurodegeneration.

The Cohen’s d effect size measures the distance between two normal distributions:

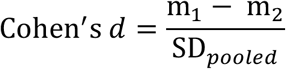

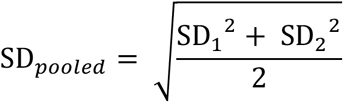

where m, and SD are the mean and standard deviation, respectively. Based on a conventional operational definition of Cohen’s *d,* small, medium and large effect sizes are defined as *d* < *0.5*, *0.5* < *d* <*0.8*, *and d* >*0.8*, respectively. We used 200 bootstrapped replicates to obtain a more robust estimation of the effect size.

Further, we used a linear regression model to estimate the association of each marker with age in each clinical group, correcting for sex and ADNI study phase. The linear regression model was fitted using the group label and age as independent variable to and corrected marker values (as dependent variables. In other words, changes in each marker value are modeled using age and the study group of the subject. Therefore, this experiment shows how each marker changes versus age in each clinical group. This can be considered as a comparison between normal and abnormal aging.

## Results

### Cohen’s d Effect size

The mean effect size (standard deviation) was 0.3415 (0.1255) for hippocampal volume, and 0.5884 (0.1215) for hippocampal SNIPE score. A pairwise t-test showed that effect sizes were significantly different between the two markers (*p* < 2.2 *e*^−16^). Hippocampal SNIPE score shows medium effect size, while hippocampal volume has a small effect size.

### Linear Regression

The graphs show hippocampal volumes (Figure 1) and SNIPE scores (Figure 2) plotted against age for both left and right hippocampi. The t-statistics and corresponding p-values show that there is a significant difference between Stables and Progressors clinical groups in either hippocampal volumes or SNIPE scores (hippocampal volumes: *t* = 2.509 (*p* = 0.012), *t* = 2.686 (*p* = 0.007; SNIPE scores: *t* = 3.222 (*P* = 0.00135) and *t* = 4.601 (*p* = 5.3*e*^−16^), for left and right respectively). There was a significant difference between Stable and Progressor groups in all experimental settings. Figure 2 shows however how this difference is emphasized in hippocampal SNIPE scores, and furthermore, is more dominant in the right hippocampus.

**Figure 1.**
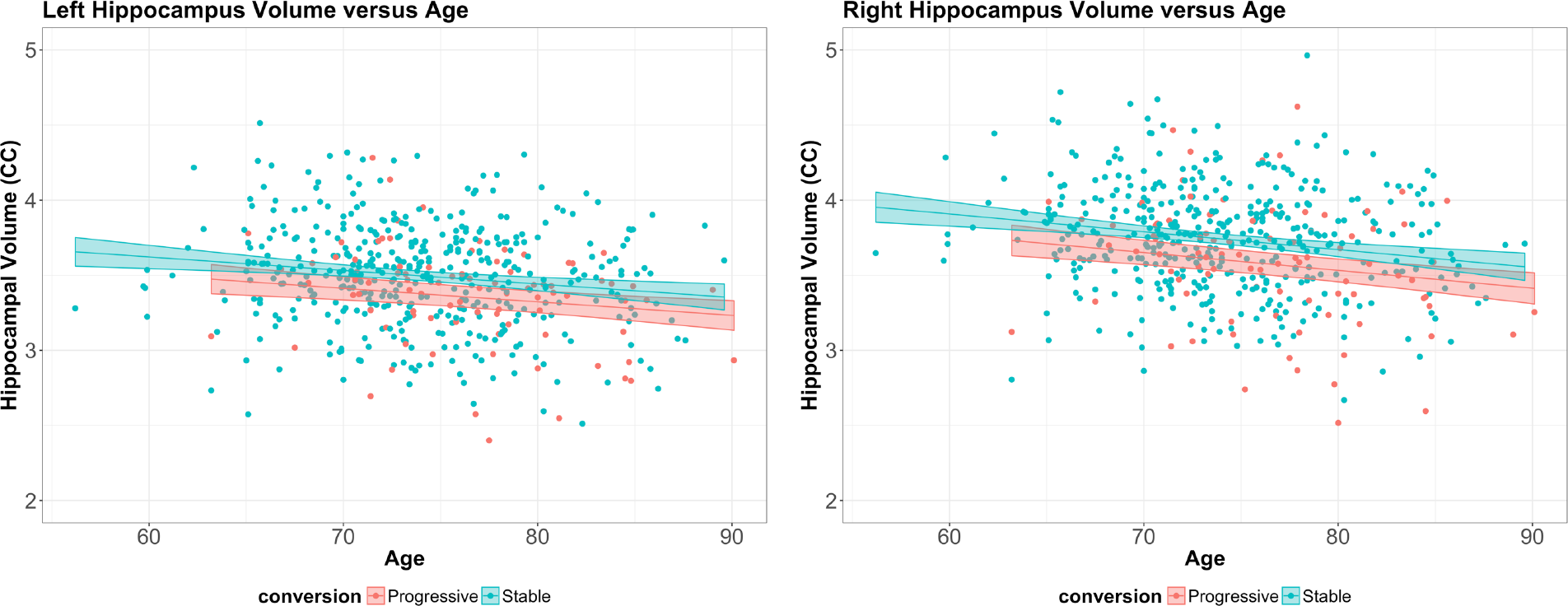
Hippocampal volume versus age for left and right hippocampus. The colors represent different clinical groups. Stable group consists the subjects who remained stable, while progressive group shows the subjects who progressed to MCI during the follow-up. The reported values are in cubic centimeters

**Figure 2.**
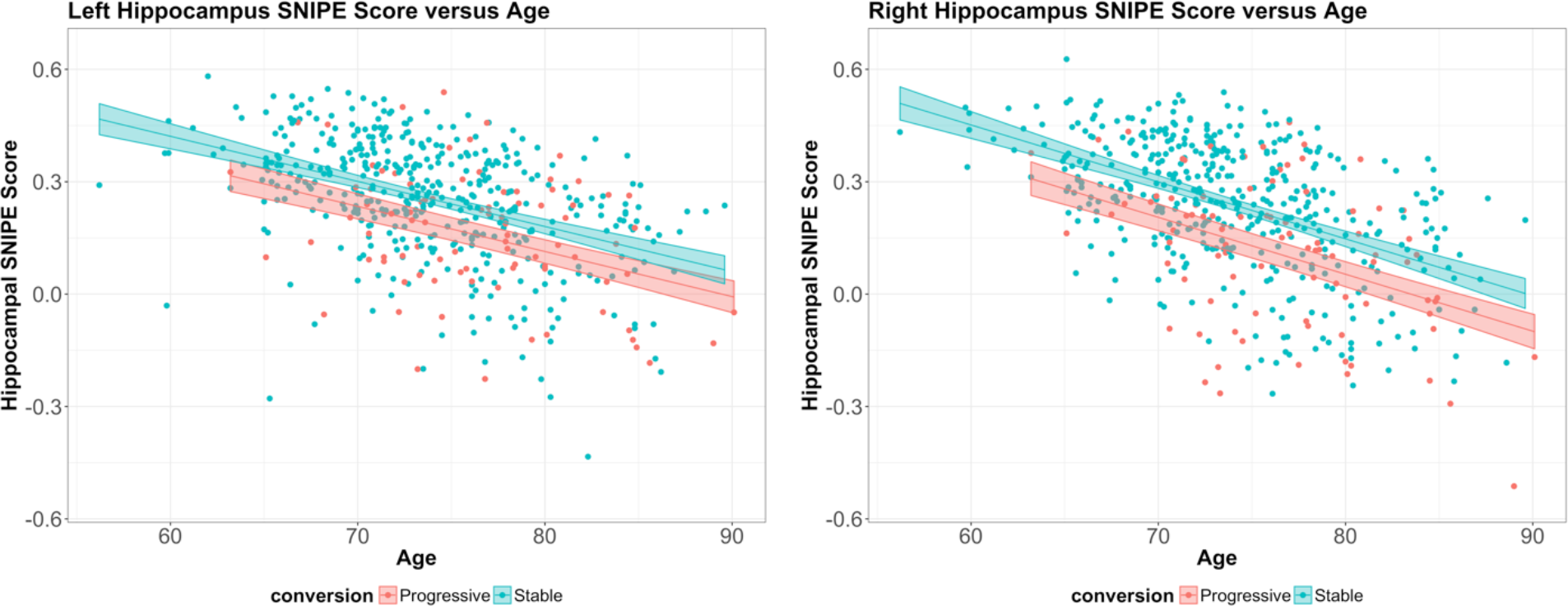
Hippocampal SNIPE score versus age for left and right hippocampus. The colors represent different clinical groups. Stable group consists the subjects who remained stable, while progressive group shows the subjects who progressed to MCI during the follow-up.

## Discussion

Early detection of Alzheimer’s disease pathology in the asymptomatic aging population may increase the effectiveness of interventional procedures to delay dementia onset (Ngandu et al., 2015). In this study we demonstrated that both hippocampi are affected even in pre-clinical stage, when the subject is considered cognitively healthy. These results are in line with the hypothesis behind the Jack biomarker model, which shows neurodegenerative atrophy, which can be captured by structural MRI, occurs before clinical symptom appear (Clifford R. Jack et al., 2010). Our effect size analyses show that Cohen’s *d* for between-group differences using hippocampal volume is small during this preclinical stage, especially when compared to the large effect size between AD patients and age-matched CH subjects (A. Zandifar et al., 2017). This suggests that to better capture early disease impact, there is a need to use more sophisticated feature extraction techniques such as our SNIPE marker, or hippocampal texture (Coupé, Eskildsen, Manjón, Fonov, & Collins, 2012; Sorensen et al., 2016). In this study hippocampal SNIPE scoring showed medium effect sizes between Stable and Progressor groups, demonstrating a higher sensitive to early AD-related changes than hippocampal volume. This observation is consistent with our previous study in MCI population where a predictive model using hippocampal SNIPE scores led to higher accuracy when compared to using hippocampal volume (Coupé, Eskildsen, Manjón, Fonov, & Collins, 2012; Coupe et al., 2015).

In order to show how ageing affects the markers in each group, we plotted the markers against age for both Stable and Progressor groups. Our result shows that there is a significant difference between the Stable group and the Progressor considering both markers, and this difference is larger for SNIPE score. However, our data did not demonstrate a different slope over time for each group. This suggests that the disease process may be delayed in the Stable group.

We also noticed that the difference between the groups in the linear regression is larger for the right hippocampus. A two-way ANOVA with hemisphere and marker (hippocampus volume and SNIPE score) as independent variables shows that the difference is significant both between the hemispheres and the markers. However, since previous studies have shown no significant difference for hippocampal hemisphere in AD detection (C. R. Jack et al., 1997), or some even showed the difference is more emphasized on the left side (Shi, Liu, Zhou, Yu, & Jiang, 2009), we believe that the observation could be specific to this particular dataset. Therefore, we decided to use mean hippocampal volume averaged over both hemispheres as a more robust marker to measure the effect size.

This paper shows that SNIPE score can even be used as an informative feature in a model to predict future amnestic MCI and future onset of dementia during the preclinical stage. This marker with more validation will probably be able to provide the clinic with a predictive tool that could have a very long prediction time margin, and clinical trials for cohort enrichment strategies.

## Acknowledgements

Data collection and sharing for this project was funded by the Alzheimer’s Disease Neuroimaging Initiative (ADNI) (National Institutes of Health Grant U01 AG024904) and DOD ADNI (Department of Defense award number W81XWH-12-2-0012). ADNI is funded by the National Institute on Aging, the National Institute of Biomedical Imaging and Bioengineering, and through generous contributions from the following: AbbVie, Alzheimer’s Association; Alzheimer’s Drug Discovery Foundation; Araclon Biotech; BioClinica, Inc.; Biogen; Bristol-Myers Squibb Company; CereSpir, Inc.; Cogstate; Eisai Inc.; Elan Pharmaceuticals, Inc.; Eli Lilly and Company; EuroImmun; F. Hoffmann-La Roche Ltd and its affiliated company Genentech, Inc.; Fujirebio; GE Healthcare; IXICO Ltd.; Janssen Alzheimer Immunotherapy Research & Development, LLC.; Johnson & Johnson Pharmaceutical Research & Development LLC.; Lumosity; Lundbeck; Merck & Co., Inc.; Meso Scale Diagnostics, LLC.; NeuroRx Research; Neurotrack Technologies; Novartis Pharmaceuticals Corporation; Pfizer Inc.; Piramal Imaging; Servier; Takeda Pharmaceutical Company; and Transition Therapeutics. The Canadian Institutes of Health Research is providing funds to support ADNI clinical sites in Canada. Private sector contributions are facilitated by the Foundation for the National Institutes of Health (http://www.fnih.org). The grantee organization is the Northern California Institute for Research and Education, and the study is coordinated by the Alzheimer’s Therapeutic Research Institute at the University of Southern California. ADNI data are disseminated by the Laboratory for Neuro Imaging at the University of Southern California.

